# Dose optimisation of favipiravir against RNA viruses using physiologically-based pharmacokinetic modelling

**DOI:** 10.64898/2026.06.24.733995

**Authors:** William Wood, Henry Pertinez, Tim Rowland, Andrew Owen, Tom Fletcher, Saye Khoo

## Abstract

Favipiravir (FPV) is an RdRp inhibitor developed and licensed in Japan for influenza but which has shown promising in-vitro activity against a range of RNA viruses. A physiologically-based pharmacokinetic model was developed for oral FPV and its metabolite M1 in order to optimise the dose regimen against plasma concentration targets for a number of viral pathogens. The model was validated using clinical data and was able to capture the variability in plasma concentrations for a population of individuals. FPV doses predicted to cause *in-vivo* exposures exceeding *in-vitro* IC_90_ targets against influenza, Ebola, Lassa fever, CCHF, SFTS, Andes virus and SARS-CoV-2, lie within the window of observed safe dosing, with SARS-CoV-2 requiring predicted doses of 2400 mg twice daily due to lower in-vitro potency. Simulations showed that a loading dose on day one of treatment should allow plasma targets to be exceeded on day one. Simulations of chronic kidney disease (CKD) showed no change to FPV plasma concentration in individuals with CKD3 and CKD5 compared to healthy individuals.

Clinical data suggested active renal efflux of M1 which led to a predicted 2.2 and 11.5 fold increase in the maximum plasma concentrations of M1 in individuals with CKD3 and CKD5 respectively in comparison with healthy individuals.

## Introduction

The COVID-19 pandemic has highlighted a need for better preparation against future outbreaks. The 100 Days Mission aims to have effective therapeutics within 100 days of the onset of a novel future pandemic. In order to meet this challenge, a library of candidate antiviral therapeutics which are ready for phase II clinical trials is required, along with a need to establish safe and effective dosing across diverse populations for viral pathogens of pandemic potential. The optimal dosage to achieve effective plasma concentrations of candidate therapeutics, either for use in phase II clinical trials or for urgent compassionate use, remains unclear for a number of viruses. In addition, there are limited pharmacokinetic data for special populations usually excluded from early phase clinical trials such as patients with chronic kidney disease (CKD).

Favipiravir (FPV) is an antiviral, originally developed as a treatment for influenza^1,2^, that has since shown *in-vitro* activity (*in-vitro* IC90 determined in Vero-E6 cells) against a wide range of viral diseases^3^ including SARS-CoV-2^3,4^, Ebola^5^, Crimean-Congo Hemorrhagic Fever (CCHF)^6^, Severe fever with thrombocytopenia syndrome (SFTS)^7,8^ and Andes virus^9^. FPV has also shown potential activity against Lassa Fever Virus in animal models^10,11^, and has recently been used in a phase II clinical trial^12^

FPV is a nucleic acid analogue prodrug which is converted intracellularly into its active form Favipiravir-ribofuranosyl-5’-triphosphate (FRTP) (Fig. 1). FRTP subsequently binds to and inhibits RNA-dependent RNA polymerase (RdRp). Clinical trials have shown that intravenous FPV is well tolerated up to 2400 mg b.d^13^ and *in-vitro* studies have shown it has a high barrier to resistance^14^. FPV exhibits wide variability in pharmacokinetics (PK) between individuals^3,15,16^ and populations^17^ and is metabolised by aldehyde oxidase (AO) (and to a lesser extent by xanthine oxidase (XO))^16^ to form the inactive metabolite M1, which is renally cleared. Variability has been observed in the expression levels^18^ and genotype^19^ of AO.

**Figure 1.**
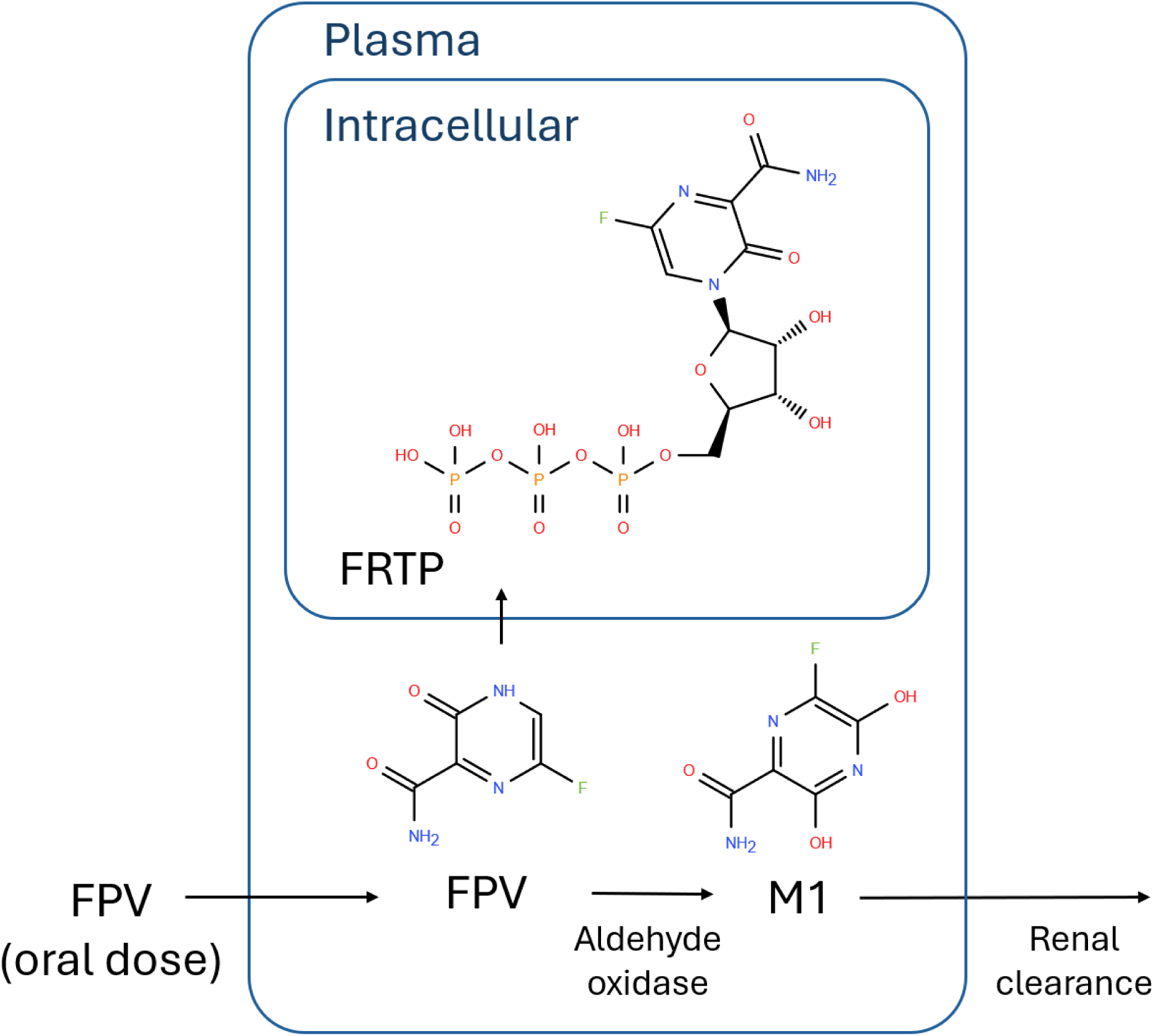
Pathway diagram for oral FPV and its metabolites FRTP and M1.

Physiologically-based pharmacokinetic (PBPK) modelling is a mechanistic mathematical modelling approach which uses the physicochemical properties of a compound and human physiological characteristics to predict the concentration-time profile of a compound in plasma and/or other required tissues. This approach can be used to establish effective dose regimens for repurposed drugs against novel pathogens using *in-vitro* efficacy (such as the IC_50_ or IC_90_) against the pathogen to estimate *in-vivo* target concentrations^20^. For antivirals, particularly for the treatment of viruses with high viral loads, fast replication rates, and/or high propensity to develop resistance, the goal is to calculate a sufficient dose such that the trough concentration at steady state at the very least exceeds the in-vitro IC90^21^, although targets for clinical efficacy may exceed this level by several multiples.

FPV is currently licensed in Japan for the treatment of influenza and the standard regimen is 1600 mg b.d. on day 1 followed by 600 mg (days 2 - 5), although clinical trials often opt for a dose regimen of 1800 mg b.d. on day 1 followed by 800 mg thereafter^1,3,4^. It is unclear however what the optimal dosage regimen should be for the treatment of other viruses. In this study, we developed and validated a PBPK model of FPV and M1 to support optimal dosing for a range of viral diseases in healthy individuals and those with CKD. Any toxicity associated with FPV as a consequence of CKD is likely due to the accumulation of M1^22^.

## Methods

A PBPK parent-metabolite model for FPV and M1 was implemented in PK-Sim (version 11.3) with input parameters as in table 1. For the purposes of this study Aldehyde oxidase, responsible for the conversion of FPV to M1, was assumed to be the only route of metabolism for FPV and renal clearance was assumed to be the only route of elimination of M1. A Michaelis-Menten model with average human observed *in-vitro* Km and V_max_ values^16^ was used for this clearance pathway. The *in-vitro* V_max_ [μM/min] was converted into the in-vivo K_cat_ [1/min] in PK-Sim by dividing by the *in-vitro* enzyme concentration, which for commercially available human liver cytosol is approximately 0.6 μM^23^. The tissue specific V_max_ is calculated in PK-Sim by multiplying the K_cat_ by the tissue AO (gene AOX1) concentration. The tissue expression of AOX1 was obtained using real-time PCR data within the PK-Sim expression database^24^. The concentration of AO in adult human hepatocytes has been measured at 10.2 pmol per million hepatocytes^18^. This was converted to 3 μM using an estimated volume of 3.4x10^-6^ L per million hepatocytes and was used as the reference concentration of AO in the liver this study.

**Table 1.** PBPK model parameters for FPV and M1.

|  | Parameter | FPV | Reference | M1 | Reference |
| --- | --- | --- | --- | --- | --- |
| Physico-chemical properties | Log P | 0.7 | <sup>35</sup> | 0.54 | DrugBank |
|  | Molecular weight (g/mol) | 157.1 | PubChem | 173.1 | DrugBank |
|  | Fraction unbound in plasma | 0.46 | Drug Label | 0.99 | Parameter identification |
|  | pKa | 5.1 (acid) | <sup>38</sup> | 9.1 | DrugBank |
|  | Aqueous solubility (g/L) | 8.5 | <sup>26</sup> | 6.7 |  |
| Physiological properties | Blood to plasma ratio | 1 | Assumed | - |  |
| | Intestinal permeability (nm/s) | $8.1 \times 10^{-4}$ cm/min | <sup>26</sup> | - | |
|  | Renal clearance (GFR fraction) | 0 | Assumed | 5.14 | Parameter identification |
| Aldehyde oxidase metabolism | $V_{\max}$ (μM/min) | 0.22 (initial), 3.21 (parameter identification) | <sup>16</sup> | / | / |
| | $K_m$ (μmol/L) | 657.5 | <sup>16</sup> | / | / |
| | Liver enzyme concentration (μmol/L) | 3 | <sup>18</sup> | / | / |
| Weibull dissolution parameters | Dissolution time (min) | 1 | <sup>25</sup> | / | / |
|  | Lag time (min) | 0 | <sup>25</sup> | / | / |
|  | Dissolution shape | 0.3 | <sup>25</sup> | / | / |

The dissolution of oral FPV was described by a Weibull function based on empirical in-vitro data^25^. The tissue specific cellular permeabilities and the tissue/plasma partition coefficients were calculated using the PK-Sim Standard and Rodgers and Rowland models respectively. For absorption, an empirical apparent permeability (based on *in-vitro* Caco-2 measurements) was used^26^

To predict the variability of plasma concentrations a population of 100 virtual individuals was created using PK-Sim. The population represented 50% females, aged 20-60 years, with bodyweights in the range 50-120 kg^21^. The variability of AO expression, represented by a standard deviation of 19.6% of mean value^18^, was also incorporated into the population.

Parameter optimisation was used to estimate the renal clearance and fraction unbound in plasma of M1 and adjust the V_max_ of AO against FPV. This was done using the Monte Carlo algorithm within parameter identification functionality in PK-Sim and used published FPV and M1 clinical data^3^. The clinical data was obtained using the Plot Digitizer online tool (plotdigitizer.com, accessed December 2025). The US316 dataset was used which contains 301 individuals in the FPV treatment arm with an average age of 41.3 years, an average bodyweight of 83.2 kg, a sex representation of 58.5% female, and who were given 1800 mg twice daily on day 1 of treatment followed by 800 mg twice daily for a further 4 days for the treatment of influenza. The lower limit of quantification (LLQ) was 0.02 μg/ml for both FPV and M1^3^.

CKD affects multiple parameters in the PK-Sim PBPK model, including kidney volume and blood flow, haematocrit and plasma protein binding^27^. For individuals with CKD3 and CKD5 we used physiological parameters provided in the ESQlabs (ESQlabs GmbH, Germany) renal impairment online course (learn.esqlabs.com/catalog/info/id:248, accessed December 2025). A table of physiological parameters is provided in table 2. For the purposes of this study, it was assumed that individuals with CKD3 and CKD5 had no change in aldehyde oxidase tissue expression or the tissue expression of any unidentified M1 transporters compared to healthy individuals.

**Table 2.** PBPK model parameters for chronic kidney disease modelling.

| Parameter | Control | CKD3 | CKD5 |
| --- | --- | --- | --- |
| Population | European | - | - |
| Gender | Male | - | - |
| Age (years) | 30 | - | - |
| Weight (kg) | 70 | - | - |
| BMI (kg/m<sup>2</sup>) | 22.6 | - | - |
| Specific blood flow rate, kidney<br>(ml/min/100g organ ) | 302.71 | 186.466 | 44.194 |
| Ontogeny factor (albumin) | 1 | 0.953 | 0.84 |
| Ontogeny factor (a1-acid<br>glycoprotein) | 1 | 1.137 | 2.062 |
| GFR (specific)<br>(ml/min/100g organ) | 26.6 | 11.406 | 1.544 |
| Hematocrit | 0.47 | 0.423 | 0.294 |
| Kidney volume (L) | 0.44 | 0.377 | 0.259 |

## Results

### Validation of the PBPK model

Simulated concentration-time profiles of FPV and M1 were compared to those of observed clinical data^3^ under the 1800 mg twice on day 1, then 800 mg twice daily regimen (fig. 2a,b). Initially the renal clearance of M1 was assumed to be equal to the glomerular filtration rate (GFR) but refined using the parameter identification function in PK-Sim, leading to an increase from the assumed 1 times GFR to 5.14 times GFR. A clearance rate of around 5 times GFR is commonly observed for tubular excreted drugs^28^ A Monte Carlo simulation of 100 virtual participants was performed to estimate the variability of FPV and M1 plasma concentrations. The interquartile range of the simulations agreed with observed data but simulations were unable to capture the lower range of FPV (0.2 μg/ml LLQ^3^) and the higher range of M1 (>20 μg/ml).

**Figure 2.**
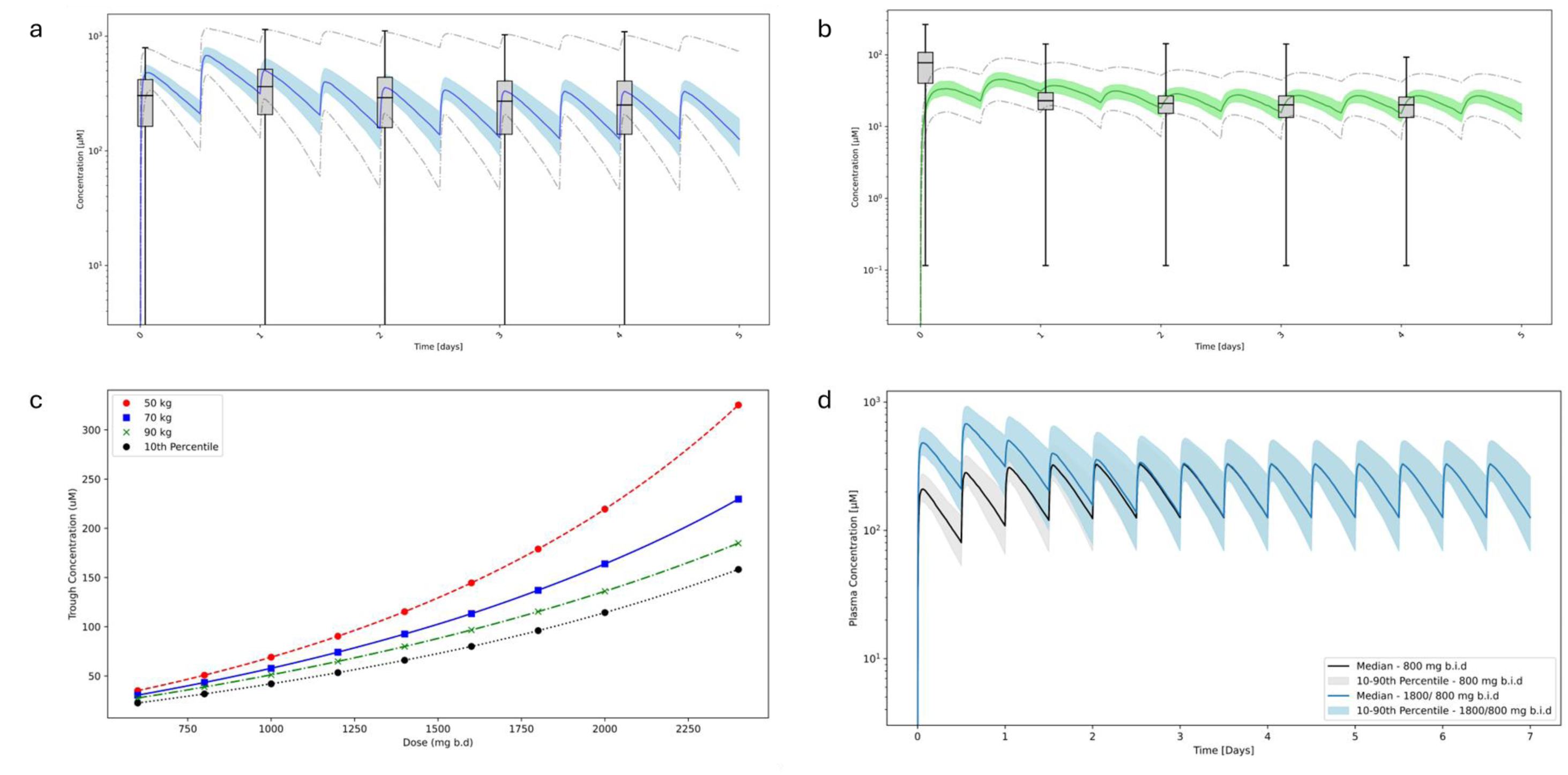
PBPK modelling of plasma concentrations of (a) FPV and (b) M1 following oral FPV dosing at 1800 mg twice daily on day 1, then 800 mg twice daily for 4 further days. a. blue solid line represents the median predicted plasma FPV concentration of an n= 100 virtual population. Blue dashed lines and grey dashed lines represent the interquartile range and the range respectively. Box and whisker plots show observed interquartile range and range of observed clinical data^3^ following the same dose regimen. b. Green solid line represents the median predicted plasma M1 concentration of an n=100 virtual population. Green dashed lines and grey dashed lines represent the interquartile range and the range respectively. Box and whisker plots show clinical data^3^. c. Dose versus unbound plasma trough concentration for 50 kg, 70 kg, and 90 kg individuals (red, blue and green markers respectively). Black circles show the 10^th^ percentile of a population simulation of 100 individuals. Lines show a spline interpolation through the points of the same colour. d. population simulations of plasma concentration versus time for an oral dose regimen of 800 mg twice daily (grey) and 1800 mg twice daily (day 1) followed by 800 mg b.d (days 2-7) (blue) to illustrate loading dose effect. Shaded areas represent the 10^th^ – 90^th^ percentiles of a simulation of 100 virtual participants.

### Dose Optimization and Loading Dose

To establish the optimal dose of FPV against a number of viruses, trough concentration-time profiles were simulated for oral FPV doses ranging from 600 mg up to 2400 mg twice daily. The steady state (day 7) trough concentrations were simulated for individuals of bodyweight 50, 70 and 90 kg (normalised to maintain a BMI of 22.6, the default value in PK-Sim) (fig. 2c). The relationship between oral dose and trough concentration was nonlinear (convex). This may be due to saturation effects as plasma concentrations of FPV in standard dose regimens such as 800 mg b.d. are comparable to the K_m_ of AO against FPV. In this study, a population simulation of 100 virtual participants was simulated, including the inter-individual variation of aldehyde oxidase expression, and the 10^th^ percentile of the trough concentration was used for dose optimisation in keeping with practice for dose optimisation of antivirals in previous studies^21^.

The optimal doses which result in predicted trough concentrations equal to the relevant IC_90_ is given for a number of viral pathogens of concern in table 3. Where the predicted dose required is less than 200 mg b.d, < 200 mg is given (200 mg is the dose in a single FPV tablet). There is still some disagreement on what is an appropriate in-vitro target for antivirals with some recommending IC_50_^20^ or multiples (4X) of IC_50_ while others have used IC_90_^21^ and we have chosen the latter as a higher level of stringency.

**Table 3.**
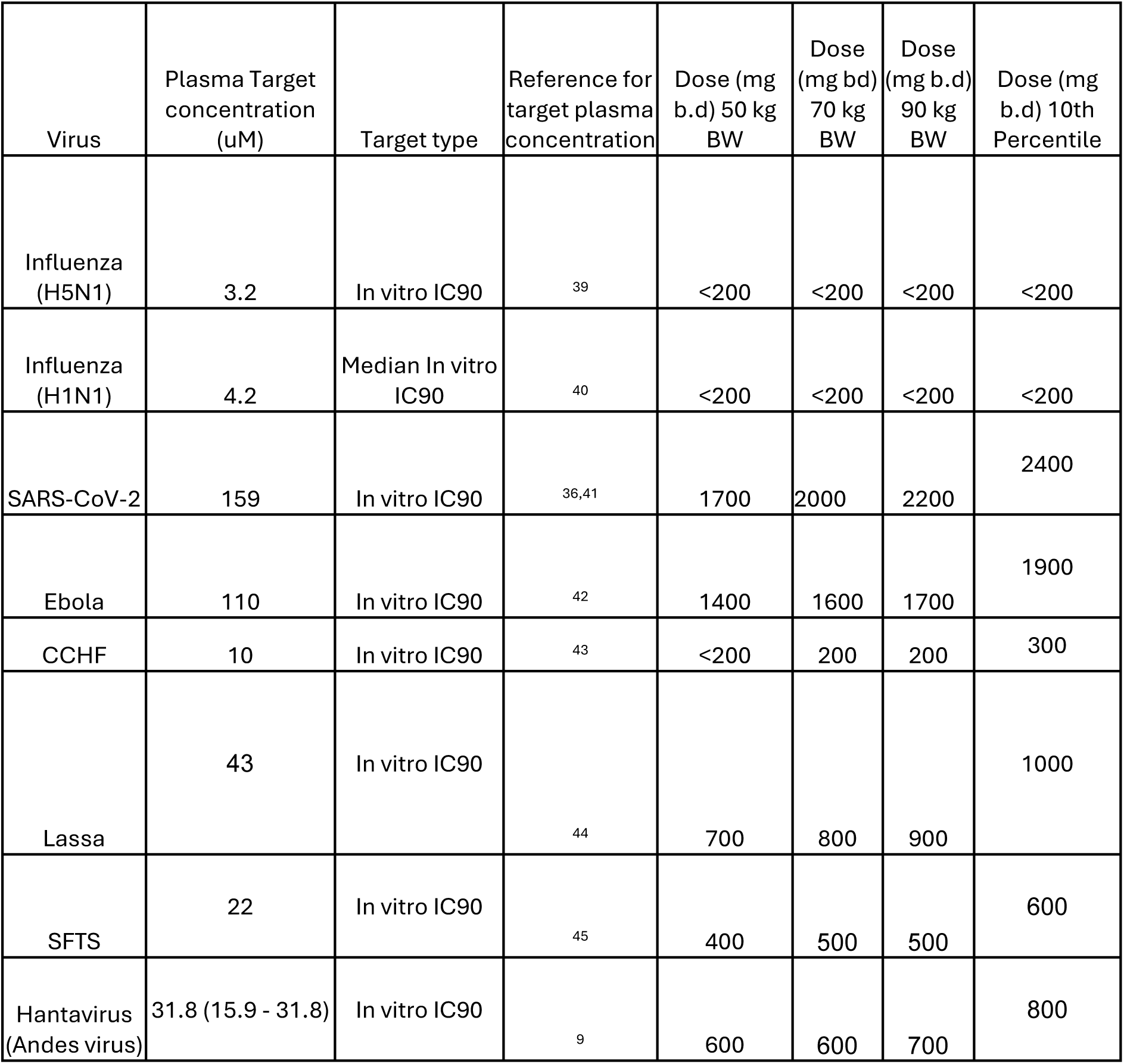
FPV Target Plasma Concentrations against various viruses. Doses are rounded to the nearest 100 mg.

With antivirals it is often important to reach efficacious plasma concentrations as soon as possible. With FPV, a loading dose is often given on day 1 (for example, 1800 mg twice on day 1, 800 mg twice daily thereafter). Simulations with and without a loading dose of 2 times the continuing dose (fig. 2d) confirm that a loading dose is necessary for plasma concentrations to exceed, and stay above, the steady state trough on day 1 of treatment.

### Renal Impairment

The impact of renal impairment on FPV and M1 plasma concentrations under standard dosing regimen was investigated within the model framework. With the PBPK model as described above used as a control, models for individuals with CKD3 and CKD5 were developed with adjusted parameters as given in table 2, with no modification of the renal clearance factor (5.14x), this therefore assumes no changes under CKD in the activity of currently unspecified transporters underlying renal clearance efflux of M1. Plasma concentration of FPV was unchanged in individuals with CKD3 and CKD5 compared to control (fig. 3). For M1 the Cmax increased from 39.55 μM in control individuals to 64.82 μM in CKD3 individuals and 221.92 μM in CKD5 individuals.

**Figure 3.**
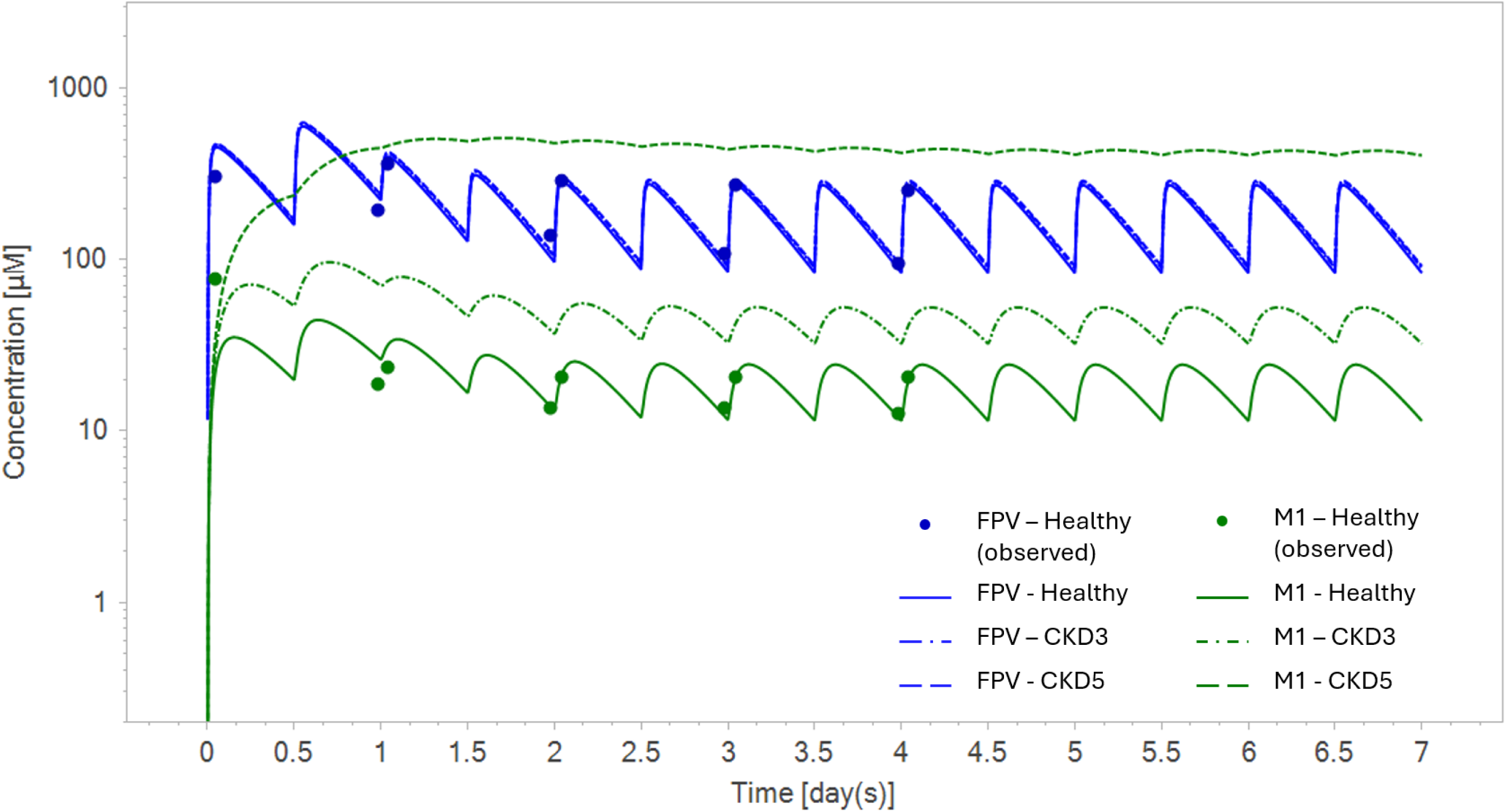
PBPK modelling of FPV and M1 in healthy individuals and individuals with chronic kidney disease (CKD). Observed plasma concentrations of FPV (blue circles) and M1 (green, circles) following an oral dose regimen of 1800 mg twice daily (day 1) and 800 mg b.d (days 2-7) in healthy individuals^3^. Overlaid with PBPK model simulations for FPV (blue) and M1 (green) for healthy individuals (solid lines), CKD3 (dash-dot lines) and CKD5 (dashed lines).

## Discussion

### Validation of the PBPK model

The PBPK model developed here showed good agreement with clinical data with adjustment of aldehyde oxidase within reasonable levels, and ability to describe the nonlinear relationship between FPV dose and plasma concentrations^29^. Plasma concentrations of FPV are highly variable. Much of this variability was captured by incorporating the variability in the expression of aldehyde oxidase into the virtual population. However, the simulations were unable to capture the lower range of FPV (0.02 μg/ml) and the higher range of M1 (>20 μg/ml) in the plasma. This may be due to the presence of individuals with a higher than usual FPV metabolism as single-nucleotide polymorphisms of human aldehyde oxidase have been shown to have a significant impact on its catalytic efficiency against a number of compounds with up to 4-fold increases in k_cat_/k_m_ observed compared to wild-type^19^. However, a simulation with a four-fold higher AO V_max_ still did not capture the lower range of FPV and higher range of M1 in this case (figure 4a,b). The model also extrapolates poorly to a very low doses. The model was further validated against observed data at a lower single 200 mg oral dose, where an increase of the V_max_ of aldehyde oxidase for the conversion of FPV to M1 by approximately 2.7 times the than the optimal V_max_ for an 800 mg twice daily dose (with a 1800 mg twice daily loading dose on day 1) was required in order for the model predictions to agree acceptably with observed data at this lower dose^30^ (fig. 4c). This may be due to minor metabolic pathways such as XO having a greater impact at very low exposures.

**Figure 4.**
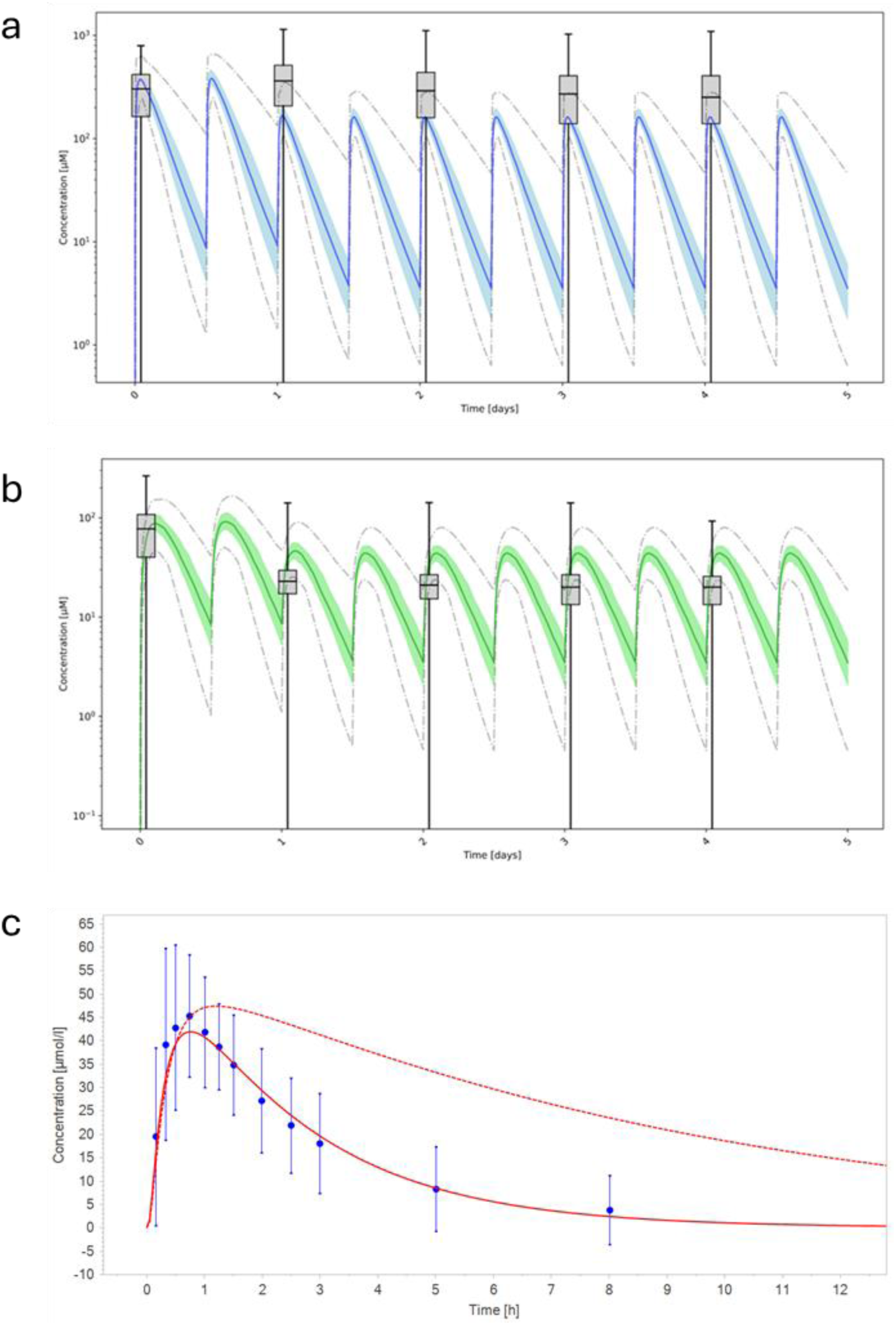
PBPK modelling of oral plasma concentrations of (a) FPV and (b) M1 following oral FPV dosing at 1800 mg twice daily on day 1, then 800 mg twice daily for 4 further days with AO V_max_ increased 4-fold to 12.84 μM/min. Oral plasma concentrations of FPV following a single oral 200 mg dose (c). a. blue solid line represents the median predicted plasma FPV concentration of a virtual population. Blue dashed lines and grey dashed lines represent the interquartile range and the range respectively. Box and whisker plots show clinical data^3^. b. Green solid line represents the median predicted plasma M1 concentration of a virtual population. Green dashed lines and grey dashed lines represent the interquartile range and the range respectively. Box and whisker plots show clinical data from the US316 clinical trial^3^. c. Red solid line represents the predicted plasma concentration of FPV following optimisation of the AO V_max_ (8.77 μM/min) on clinical data^30^ (blue dots and error bars show mean and standard deviation respectively). Red dashed line represents the predicted plasma concentration of FPV from the model optimised for the standard dose regimen which has an AO V_max_ of 3.21 μM/min.

Due to the lack of tissue expression data for xanthine oxidase in PK-Sim, the model was unable to incorporate this route of metabolism. This seems to have negligible impact on the performance of the model against clinical data. Because XO appears to contribute more at low exposures, and the lowest predicted doses (influenza, Lassa, <200 mg b.d.) fall below the calibrated range, confidence in these specific predictions is reduced; this is the dose range where omission of XO is most likely to matter. The renal clearance of M1 at 5.14 times GFR suggests there is tubular secretion, most likely via an active efflux transporter which remains to be characterised. The observed plasma concentration of M1 shows a peak on day 1 which was not predicted by the model. This may be due to autoinhibition of aldehyde oxidase by FPV.

### Dose Optimization and Loading Dose

The optimal dose for oral FPV given twice a day required to reach plasma concentrations exceeding in-vitro IC_90_ in 90% of the virtual population was predicted for a number of viruses. Low doses were required for influenza A virus and CCHF at 200 mg and 300 mg respectively twice daily. For SFTS, Andes virus and Lassa virus slightly higher doses of 600 mg, 800 mg and 1000 mg b.d respectively were required. Much higher doses were required for the treatment of Ebola virus and SARS-CoV-2: 1900 mg, and 2400 mg b.d respectively. Given FPV only shows modest clinical efficacy against SARS-CoV-2 with the standard dose of 1800 mg twice daily day 1 and 800 mg b.d. thereafter^4^, these simulations are in keeping with recommendations that the dose may need to be increased, which is also in agreement with the high dose requirement for FPV against SARS-CoV-2 observed in hamster models^31^. These are dose predictions intended to support, rather than replace, clinical evaluation. In the PIONEER trial reaching this target was associated with numerically but not significantly improved outcomes for COVID-19^32^ while intravenous dosing reached pre-specified targets without a clear efficacy signal but this was a hospitalised cohort of frail, elderly patients with moderate to severe disease^13^. Previous modelling studies have also highlighted the need for a higher dose in the treatment of Ebola virus disease with a suggested regimen of 6000 mg day 1 and 1200 mg twice daily for the following 10 days^33^.

FPV has been shown to be safe at a dose of 2400 mg b.d^12,13^, which is higher than (or equal to in the case of SARS-CoV-2) the predicted required dose for each of the viruses included in this study which suggests that FPV is a potential candidate therapy for a range of pathogenic viruses. The model described here was calibrated to approved dosing (800 mg twice daily preceded by a loading dose on Day 1) and given the non-linear dose-concentration relationship for FPV less confidence can be attached to predicted concentrations from lower doses since PK data are sparse. Additionally, the half-life of FPV is higher after multiple doses than that of a single dose^29^. For a single 200 mg dose, the half-life of FPV was 1.7 h^30^. For a range of single doses (30 – 2400 mg) FPV has been found to have a half-life of 2 - 5.5 h^29^. This model predicts a 6.1 h half-life for a multiple dose regimen of 1800 mg (day 1) 800 mg (days 2 – 14) twice daily This longer half-life may be in part due to accumulation of FRTP over multiple doses. Caution should be exercised however as a higher dose than predicted using this model may be needed if clearance is dramatically increased at lower doses beyond that predicted by the *in-vitro* Michaelis-Menten kinetics of AO against FPV.

It should be noted that the *in-vitro* targets used in this study were not corrected for *in-vitro* protein binding due to lack of data. Free drug theory holds that potency should be compared on a free-versus-free basis, but the *in-vitro* activity data were generated in serum-containing media and so the nominal IC_90_ already reflects some degree of binding; correcting only the plasma concentration to its unbound fraction while leaving the *in-vitro* target uncorrected would overstate the dose required and risks discounting a potentially useful drug^34^. As FPV demonstrates low, predominantly albumin-mediated binding^35^, an assumption of a fraction unbound *in-vitro* of ∼1 is reasonable and the two corrections act in opposing directions, such that the uncorrected comparison used here is a fair approximation. The *in-vitro* methodologies were also inconsistent in terms of the media used (eg. the concentration of fetal bovine serum or bovine serum albumin) which may affect protein binding *in-vitro.* It should also be noted that *In-vitro* methods for establishing the potency of antivirals are not standardised. Though care was taken to ensure the targets were comparable such as using IC_90_ values from vero E6 cell lines throughout this study, the methodologies employed in the assays and subsequent analysis may differ study-to-study.

The integration of a loading dose into the regimen on day 1 of at least 2 times the maintenance dose thereafter predicted plasma concentrations to exceed (and remain above) target levels within minutes whereas without a loading dose, trough concentrations on day 1 dropped below target, with a risk therefore of compromising drug effectiveness. It is worth noting that with a loading dose, the plasma concentration decreases over the first few days until reaching steady state. This has been observed in clinical trials with a loading dose^1^ but not without a loading dose^13^. The model presented here captures the nonlinear pharmacokinetics of FPV and is able to replicate both scenarios.

In this study the plasma target concentrations were based on FPV, not intracellular levels of its active metabolite FRTP. With a loading dose, it is possible to exceed plasma targets within minutes. The intracellular concentration of FRTP however, may take up to 9 hours to reach a maximum^2^. Simulations also predict the intracellular concentration of FRTP to be much lower than the plasma concentration of FPV^36^.

### Renal Impairment

PBPK modelling of renal impairment predicted no significant change in plasma FPV concentration but a significant increase in M1 concentrations. The fold difference in Cmax for M1 in individuals with CKD3 (CKD3/control) was predicted 2.2. This is in agreement with the FPV label, which states that the trough concentration of M1 is increased 2.5 times in patients with moderate renal impairment^22,37^. For individuals with CKD5, fold difference in steady state trough concentration for M1 (CKD5/control) was predicted to be 11.5. As it has been suggested that adverse effects associated with FPV are due to M1, these results suggest that caution should be taken with patients with severe renal impairment. The fitted renal clearance of M1 was assumed to be unchanged between healthy and renal-impaired individuals. This may not be true *in vivo* if there unidentified M1 transporters which differ in expression between healthy and renal-impaired individuals. Due to lack of evidence to the contrary, it was also assumed for the purpose of this study that aldehyde oxidase expression was unchanged in renal-impaired individuals.

## Conclusion

PBPK modelling of antivirals can predict optimal dosing regimens in humans against a range of viruses when clinical data is lacking. In this study, a PBPK model was developed for the RdRp inhibitor FPV and was used to predict optimal dose regimens for FPV against influenza, SARS-CoV-2, Ebola, Lassa fever, CCHF, SFTS, and Andes virus, using *in-vitro* target concentrations. In all cases plasma targets were found to be achievable with doses known to be safe.

## Author Approvals

All authors have read and approve of this article.

